# RNA-Targeting CRISPR/Cas13d System Eliminates Disease-Related Phenotypes in Pre-clinical Models of Huntington’s Disease

**DOI:** 10.1101/2022.01.23.477417

**Authors:** Kathryn H. Morelli, Qian Wu, Maya L. Gosztyla, Hongshuai Liu, Chuangchuang Zhang, Jiaxu Chen, Ryan Marina, Kari Lee, Krysten L. Jones, Wenzhen Duan, Gene W. Yeo

**Affiliations:** Division of Neurobiology, Department of Psychiatry and Behavioral Sciences, Johns Hopkins University School of medicine, Baltimore, Maryland, USA; School of Traditional Chinese Medicine, Beijing University of Chinese Medicine, Beijing, China; Department of Cellular and Molecular Medicine, University of California San Diego, La Jolla, California, USA; Institute for Genomic Medicine, University of California San Diego, La Jolla, California, USA; Stem Cell Program, University of California San Diego, La Jolla, California, USA; The Solomon H Snyder Department of Neuroscience, Johns Hopkins University School of Medicine, Baltimore, Maryland, USA; Program in Cellular and Molecular Medicine, Johns Hopkins University School of Medicine, Baltimore, Maryland, USA

## Abstract

Huntington’s disease (HD) is a fatal, dominantly inherited neurodegenerative disorder caused by a CAG trinucleotide expansion in exon 1 of the huntingtin (*HTT*) gene. Although the pathogenesis of HD remains complex, the CAG-expanded (CAG^EX^) *HTT* mRNA and protein ultimately causes disease through a toxic gain-of-function mechanism. As the reduction of pathogenic mutant *HTT* mRNA is beneficial as a treatment, we developed a CAG^EX^ RNA-eliminating CRISPR-Cas13d system (Cas13d/CAG^EX^) that binds and eliminates toxic CAG^EX^ RNA in HD patient iPSC-derived striatal neurons. We show that intrastriatal delivery of Cas13d/CAG^EX^ via a single adeno-associated viral vector, serotype 9 (AAV9) mediates significant and selective reduction of mutant *HTT* mRNA and protein levels within the striatum of heterozygous zQ175 mice, an established mouse model of HD. Moreover, the reduction of mutant *HTT* mRNA renders a sustained reversal of HD phenotypes, including improved motor coordination, attenuated striatal atrophy, and reduction of mutant HTT protein aggregates. Importantly, phenotypic improvements were durable for at least 8 months without gross or behavioral adverse effects, and with minimal off-target interactions of Cas13d/CAG^EX^ in the mouse transcriptome. Taken together, we demonstrate a proof-of-principle of an RNA-targeting CRISPR/Cas13d system as a therapeutic approach for HD, a strategy with broad implications for the treatment of other dominantly inherited neurodegenerative disorders.

## Introduction

Huntington’s disease (HD) is a common autosomal dominant neurodegenerative disorder, caused by a CAG short tandem repeat (STR) expansion in exon 1 of the Huntingtin (*HTT*) gene (Paulson 2018). Onset of the motor symptoms, known as the clinical onset of HD, can occur from childhood to old age, with a mean age of onset at ∼45 years followed by inexorable disease progression. The therapies currently available to HD patients offer only moderate symptom relief, and the affected individuals typically die 15–20 years post-diagnosis due to complications (Langbehn et al 2010, Ross et al 2014).

Mutant HTT protein (mHTT) affects a variety of cellular functions. It binds and interacts with DNA in many genes, resulting in transcriptional dysregulation (Hensman Moss et al 2017, Hodges et al 2006), neuronal dysfunction and, eventually, degeneration. Considering the pathogenic events that occur downstream of mHTT form a complex web, targeting of individual pathways is either too difficult to achieve cleanly or insufficient to modify the disease course of HD. A major focus of HD therapeutic development has recently shifted towards targeting the root of the disease, causative mutant *HTT* (Tabrizi et al 2019, Wild & Tabrizi 2017). Besides the toxicity of the mutated HTT protein, an increasing body of evidence indicates that mutant *HTT* mRNA also contributes to disease pathogenesis (Banez-Coronel et al 2012, de Mezer et al 2011); consequently, strategies to suppress both *HTT* transcripts and protein levels would be most beneficial as a treatment. RNA interference (RNAi) and antisense oligonucleotide (ASO) strategies have shown preclinical efficacy and are being tested in clinical trials (Tabrizi et al 2019). However, most of these approaches do not precisely differentiate mutant *HTT* from the normal allele (McColgan & Tabrizi 2018).

Most HD patients are heterozygous for the CAG expansion, and normal HTT plays important roles in brain development as well as the adult central nervous system (CNS) (Saudou & Humbert 2016). In the adult brain, HTT plays a part in intracellular vesicle trafficking (Brandstaetter et al 2014, Caviston & Holzbaur 2009, Strehlow et al 2007), transcriptional regulation (Kegel et al 2002, Zuccato et al 2003), and synaptic connectivity (DiFiglia et al 1995, Marcora & Kennedy 2010, McKinstry et al 2014). While partial reduction of normal *HTT* levels is tolerable for a short period, long-term (years) ramifications of reductions are unclear given its involvement in a myriad of biological functions. Sustained reduction of normal *HTT* levels might exacerbate HD pathogenesis (Gauthier et al 2004, Saudou & Humbert 2016). The recent termination of the phase III clinical trial of an ASO (non-allele selective HTT-lowering) in HD patients underscores the urgency to develop strategies that can selectively and effectively suppress mutant *HTT* expression (Gagnon et al 2010, Garriga-Canut et al 2012, Yu et al 2012).

The RNA-guided RNA-targeting subtype of IV-D CRISPR/Cas, Cas13d, has been recently identified as an efficient and specific RNA targeting approach that can be applied in mammalian cells (Zhang et al 2018). Cas13d, the smallest known Cas protein with a size of ∼930 amino acids, can be packaged and delivered with its sgRNAs to target cells via a single AAV capsid. Cas13d possesses dual RNase activities and is capable of processing CRISPR arrays and cleaving target RNAs in a protospacer flanking sequence (PFS)-independent manner (Konermann et al 2018, Yan et al 2018, Zhang et al 2018). In cell-based screening as well as side-by-side comparisons to short-hairpin RNA, nuclear-localized sequence (NLS)-fused Cas13d showed a strong ability to cleave target RNA with high efficiency (∼96% knockdown by Cas13d comparing to ∼65% by shRNA), high specificity, and no significant off-target effects in mammalian cell culture (Konermann et al 2018). These findings indicate that CRISPR/Cas13d is a promising platform for RNA targeting. Moreover, Cas13d targets RNA, which may have fewer irreversible side effects resulting from permanent gene editing. The high specificity, significant efficiency, and ease of operation render a CRISPR/Cas13d-mediated RNA editing approach advantageous in potential clinical applications.

In this report, we tested a CRISPR/Cas13d-based gene therapy approach that silences mutant toxic CAG-expanded (CAG^EX^) *HTT* RNA in both human patient iPSC models and an established mouse model of HD. We show that our CAG^EX^-targeting Cas13d system (Cas13d/CAG^EX^) selectively reduces CAG^EX^ *HTT* RNA in striatal neurons derived from three HD patients with CAG expansions ranging from 66 to 109 short tandem repeats. AAV-mediated delivery of Cas13d/CAG^EX^ to the striatum of zQ175/+ HD mice resulted in allele-selective suppression of mutant HTT protein while maintaining normal HTT levels. Furthermore, administration of Cas13d/CAG^EX^ in premanifest zQ175/+ HD mice significantly attenuated striatal atrophy, improved motor function, and reduced mHTT aggregates in the striatum. Our data provided the first evidence that CRISPR/Cas13d mediated elimination of mutant *HTT* mRNA and protein is a promising therapeutic approach to develop further for HD treatment.

## Results

### Design and initial screening of an RNA-targeting Cas13d system that targets toxic CAG expansions

Although the etiology of HD remains complex, many proposed mechanisms arise from the transcription and subsequent translation of CAG^EX^ *HTT*, ultimately causing disease through a toxic gain-of-function mechanism. Therefore, therapeutic approaches that suppress mutant *HTT* at the RNA level are actively being pursued. We have recently pioneered the repurposing of the Cas9 system to target and eliminate toxic repeat RNAs *in vitro* (Batra et al 2017) and *in vivo* (Batra et al 2021) in a two-vector AAV system, demonstrating that RNA-targeting CRISPR approaches are particularly effective with RNA repeat expansions. Here, due to the advantageous compact nature of *Ruminococcus flavefaciens* XPD3002 (Rfx) CRISPR/Cas13d, we developed a CAG^EX^ RNA-targeting Cas13d (Cas13d/CAG^EX^) system that we packaged into a single vector in both lentiviral and adeno-associated viral delivery vehicles to evaluate its therapeutic potential in multiple established pre-clinical models of HD, such as striatal neurons from a panel of HD patient iPSC-lines and a full-length *HTT* knock-in mouse model expressing a human mutant exon-1 with the expanded CAG repeat (∼220 repeats) within the native mouse huntingtin gene, zQ175/+(Figure 1A).

**Figure 1.**
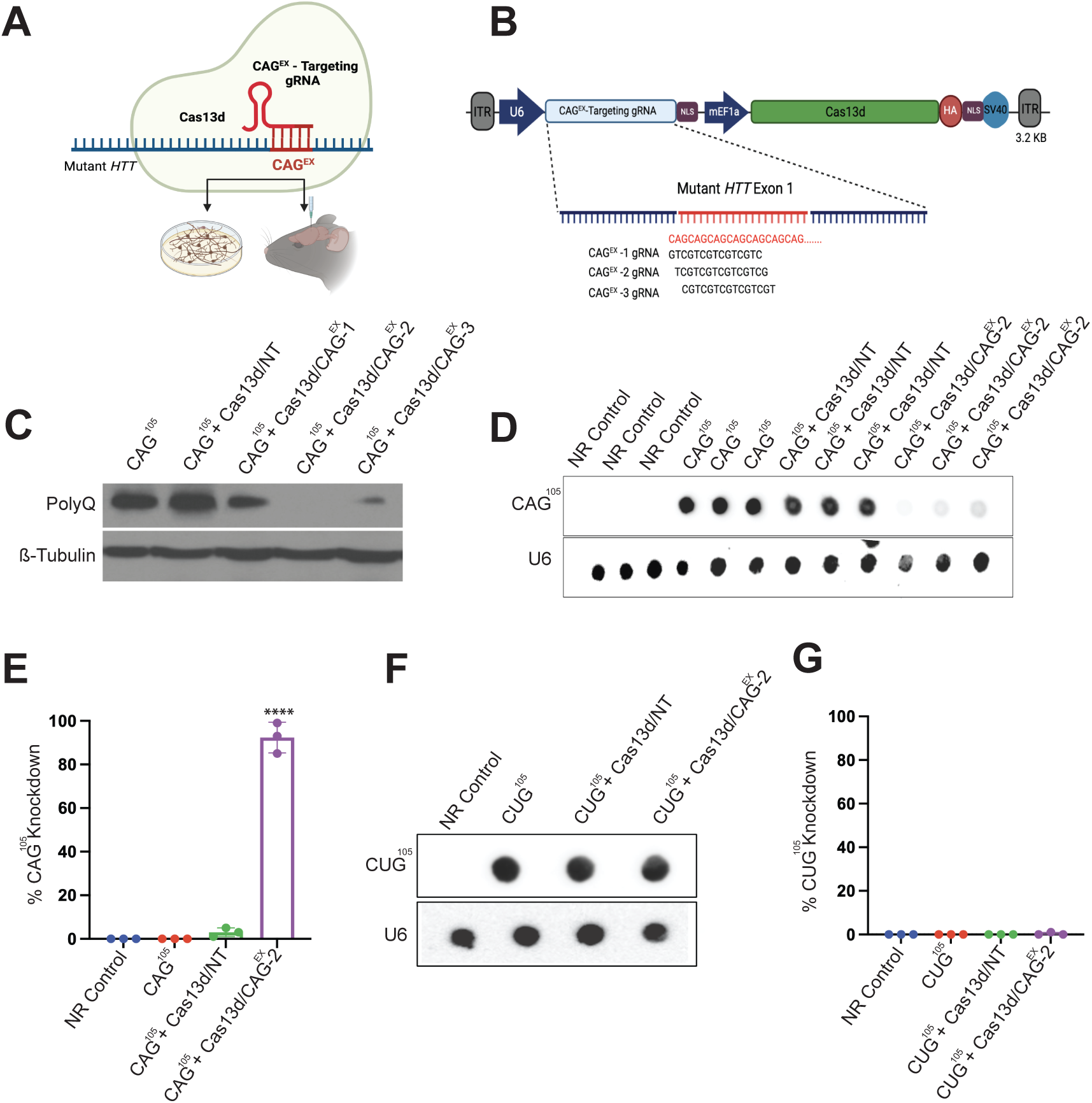
Development of an RNA-targeting Cas13d-based gene therapy approach for HD. A. Treatment scheme of our single gene therapy that expresses Cas13d and a single gRNA designed to eliminate CAG-expanded *HTT* RNA in both human striatal neuronal cultures derived from patient iPSCs and in the striatum of an established mouse model of HD, zQ175/+. B. Diagram of a series of CAG-expanded RNA-targeting vectors that consists of (1) Cas13d tagged with an HA epitope and (2) one of three U6-promoter-driven *Ruminococcus flavefaciens* XPD3002 (Rfx) CRISPR/Cas13d guide RNAs (denoted as CAG^EX^ gRNA 1-3). C. Western blot analysis of poly-Q protein from protein lysates isolated from HEK293s transfected with a CAG^105^ repeat plasmid and each candidate Cas13d vectors. D, E. RNA dot blot analysis and quantification of CAG-expanded RNA within HEK293s transfected with a CAG^105^ repeat plasmid along with (NT) or CAG^EX^1-3 vectors. F, G. RNA dot blot analysis and quantification of CUG-expanded RNA within HEK293s transfected with a CUG^105^ repeat plasmid along with a (NT) or CAG^EX^-2 vector.

First, to optimize knockdown of CAG^EX^ *HTT* RNA by Cas13d, we engineered three distinct RNA-targeting vectors that consist of (1) Cas13d tagged with an HA epitope and (2) one of three U6-promoter-driven Cas13d guide RNAs (denoted as CAG^EX^ gRNA 1-3) (Konermann et al 2018). All three gRNAs are complementary to the CAG^EX^ RNA sequence with each guide targeting a different codon within the repeat expansion: CAG^EX^ gRNA -1 (GTC), CAG^EX^ gRNA -2 (TCG), and CAG^EX^ gRNA -3 (CGT) (Figure 1B). As aggregation of toxic polyglutamine protein (poly-Q) translated from CAG^EX^ RNA is well documented as one of the primary causes of neurodegeneration in HD, we determined if and to what extent Cas13d in conjunction with each CAG^EX^-targeting gRNA can eliminate poly-Q protein in live human cells. HEK293s were co-transfected with a repeat expansion plasmid with 105 CAG short tandem repeats along with a Cas13d-containing vector with a non-targeting gRNA designed to target the bacterial transcript Lambda (Cas13d/NT) or one of the three CAG^EX^-targeting guides. Poly-Q protein in cells transfected with Cas13d and CAG^EX^ 2 was significantly reduced by 99±7%, as measured by western blot analysis (one-way ANOVA p < 0.001) (Figure 1C). RNA blots performed with RNA isolated from cells co-transfected with our CAG repeat expression plasmid and Cas13d with the CAG^EX^-2 gRNA also showed a significant 85±4% reduction of CAG^EX^ RNA compared with cells transfected with our non-targeting Cas13d vector (one-way ANOVA p < 0.001) (Figure 1D and quantified in Figure 1E). Importantly, expression studies using CUG expansion (CUG^EX^) RNA-expressing plasmids and a semi-quantitative RNA dot blot were performed to determine if Cas13d with the CAG^EX^-2 gRNA specifically eliminates RNA transcripts that contain CAG expansions without degrading other GC-rich transcripts. Excitingly, we observed that Cas13d with the CAG^EX^-2 gRNA specifically targets and degrades CAG^EX^ transcripts while leaving CUG^EX^ transcripts intact (Figure 1F, quantified in 1G). Taken together, these data confirm that our Cas13d system that includes the CAG^EX^-2 gRNA, now referred to as “Cas13d/CAG^EX^”, can effectively eliminate CAG^EX^ RNA and subsequent poly-Q protein in human cells.

### Cas13d/CAG^EX^ prevents molecular phenotypes of HD in patient iPSC-derived striatal neurons

To demonstrate the therapeutic potential of our Cas13d approach in a human preclinical model, a panel of striatal neuron cultures enriched for medium spiny neurons (MSNs) was generated from three previously validated iPSC lines derived from individual patients with HD, with CAG repeats in *HTT* ranging from 66 to 109 (HD 66, HD 77, and HD 109), and three independent unaffected neurotypical controls, denoted as Control 1-3 (Gore et al 2011, Smith-Geater et al 2020, The HD iPSC Consortium, 2012, Telezhkin et al 2016). Each HD and Control MSN line was transduced with a constitutive, lentiviral system supplying either Cas13d/CAG^EX^ or non-targeting Cas13d/NT, 16 days post-differentiation, when most of each culture consisted of neuroprogenitor cells. At day 32 post-differentiation both HD and Control cultures consisted of mature MSNs defined by cells positive for mature MSN markers, microtubule-associated protein 2 (MAP2) and Forkhead box protein P1 (FoxP1), as well as GABAergic neuronal markers, gamma aminobutyric acid (GABA) plus glutamate decarboxylase-67 (GAD-67) (Figures 2A and 2B). Widespread transduction was confirmed with Cas13d-HA-immunoflouresence with 78% or more (average 83% ± 4) of neurons showing Cas13d expression (Figures S2A-D). RNA slot blot hybridization detected CAG^EX^ RNA in all three HD lines that was significantly reduced by our Cas13d/CAG^EX^ with an 84±1% reduction in HD 66, a 79±8% reduction in HD 77, and a 56±2% reduction in HD 109 (quantified in Figure 2C, blot shown in Figure 2D).

**Figure 2.**
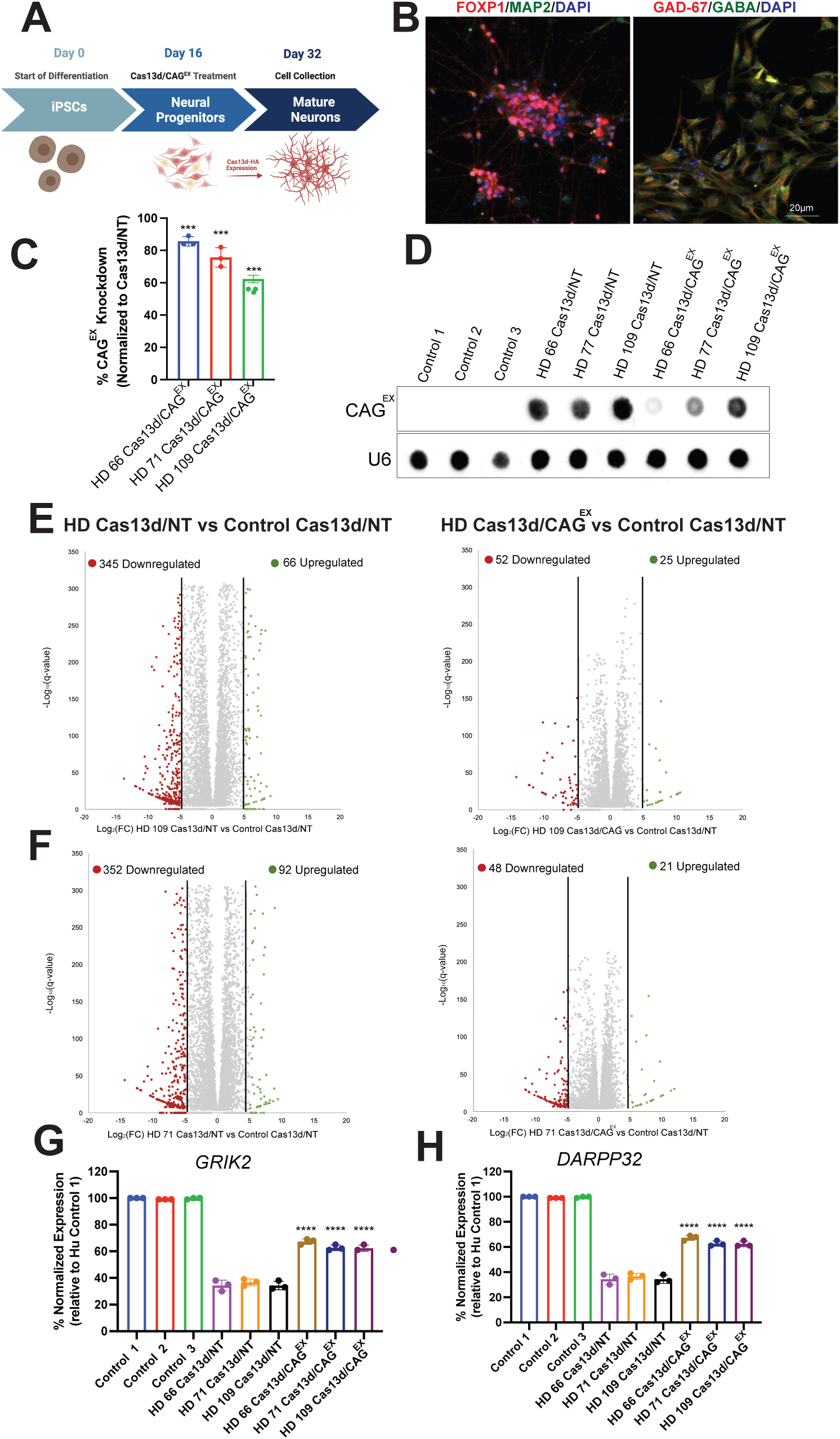
Reversal of HD-mediated molecular phenotypes within a panel of HD striatal neurons by Cas13d/CAG^EX^. A. Schematic of the differentiation protocol used for iPSC-derived striatal neurons enriched for medium spiny neurons (MSNs) treated with a lentiviral system expressing Cas13d/CAG^EX^. B. Representative immunofluorescence images of cells at day 32 in control MSNs showing that the cells are positive and co-localize for MAP2 and FOXP1 (left); cells are also positive for GABA and GAD-67 (right). Scale bar, 20 μm C, D. RNA dot blot analysis and quantification of CAG-expanded RNA within MSNs transduced with Cas13d/NT or Cas13d/CAG^EX^ –expressing lentiviral vectors. E, F. Scatter plots of upregulated and downregulated DEGs of within HD 109 and HD 77 treated with either Cas13d/NT or Cas13d/CAG^EX^ lentiviral vector showing reversal of HD-mediated changes in the human transcriptome by Cas13d/CAG^EX^ (Wilcoxon Test, p< 0.0001). G, H. Quantitation of established HD MSN marker transcripts, *GRIK2* and *DARPP32* by quantitative PCR in HD MSN cultures treated with Cas13d/NT and Cas13d/CAG^EX^.

Transcriptome-wide RNA-seq analysis was used to identify differentially expressed genes (DEGs) among all three control lines and each individual HD mature MSN culture treated with Cas13d/CAG^EX^ or Cas13d/NT. Defining DEGs at a false discovery rate of p <0.0001, we identified hundreds of down and upregulated genes in the HD lines, compared to controls (left panels in Figures 2E, 2F, Supplementary Figure. 2A, Supplementary Table 1). Excitingly, we determined that these HD-associated transcriptional changes were reversed by 84.5% in HD109 and 81.3% in HD71, and to a lesser degree (31.4%) in HD66 (that harbored ∼18% of the extent of changes in the HD71 and HD109) by Cas13d/CAG^EX^ treatment (right panels in Figures 2E, 2F, Supplementary Figure. 2B). Our results demonstrate that our gene therapy approach reduces the most prominent disease markers (Supplementary Table 1). Among these transcripts were established molecular biomarkers of HD, *GRIK2* and *DARPP32*, which were significantly upregulated by Cas13d/CAG^EX^ compared to Cas13d/NT transduced lines (Figures 2G and 2H).

Importantly, neither quantification of *HTT* mRNA by transcript expression levels (TPM) nor quantitative PCR could detect a change in total *HTT* mRNA levels in any of the three control lines treated with Cas13d/CAG^EX^ compared to Cas13d/NT or left untreated (One-Way ANOVA, Tukey’s posthoc test, p< 0.05) (Supplementary Figure. 2C, D). Significant alterations in transcripts that contain benign CAG short tandem repeats were also undetected in any of the control MSN cultures treated with Cas13d/CAG^EX^ (Supplementary Figure 2E). Taken together, these data confirm that Cas13d/CAG^EX^ can reduce toxic CAG^EX^ RNA in MSNs, and reverse HD-mediated changes in the human transcriptome without interfering with wildtype *HTT* or other transcripts with normal CAG expansions.

### Therapeutic efficacy and safety of AAV9 packaged Cas13d/CAG^EX^ in a Full-length mHTT Knock-in Mouse Model

To determine if our approach can prevent or halt the progression of neurodegenerative phenotypes in zQ175/+ mice, we packaged Cas13d/CAG^EX^ and Cas13d/NT into single stranded AAV9 viral vectors and conducted bilateral intrastriatal injection of AAV/CAG^EX^ or AAV/NT to an equal number of 2-month-old zQ175/+ HD mice and age-matched wildtype littermate controls (WT). At three weeks post injection, we observed successful expression of both Cas13d-HA fusion constructs in mouse striatum, indicated by red fluorescent signals of HA protein with an anti-HA antibody (Supplementary Figure. 3A). Motor function and body weight were monitored longitudinally over 8 months post-injection, and HTT protein levels were determined at the end of the study (Figure 3A). We evaluated the effect of treatment on motor function on a balance beam. While normally zQ175/+ HD mice exhibit clear motor deficits at 8 months of age, those treated with Cas13d/CAG^EX^ showed statistically significant improvements in motor performance, indicated by shorter traverse times, compared with zQ175/+ mice treated with Cas13d/NT (**p<0.01) or left untreated (**p<0.01) (Figure 3B). Moreover, Cas13d/CAG^EX^ injected zQ175/+ mice showed improvements in body weight compared to untreated or Cas13d/NT-treated cohorts (Figure 3D). In addition, Cas13d/CAG^EX^ had no significant effect on body weight nor motor function in WT mice (Fig 3C, E), implying that Cas13d/CAG^EX^ does not produce gross adverse effects in mice.

**Figure 3.**
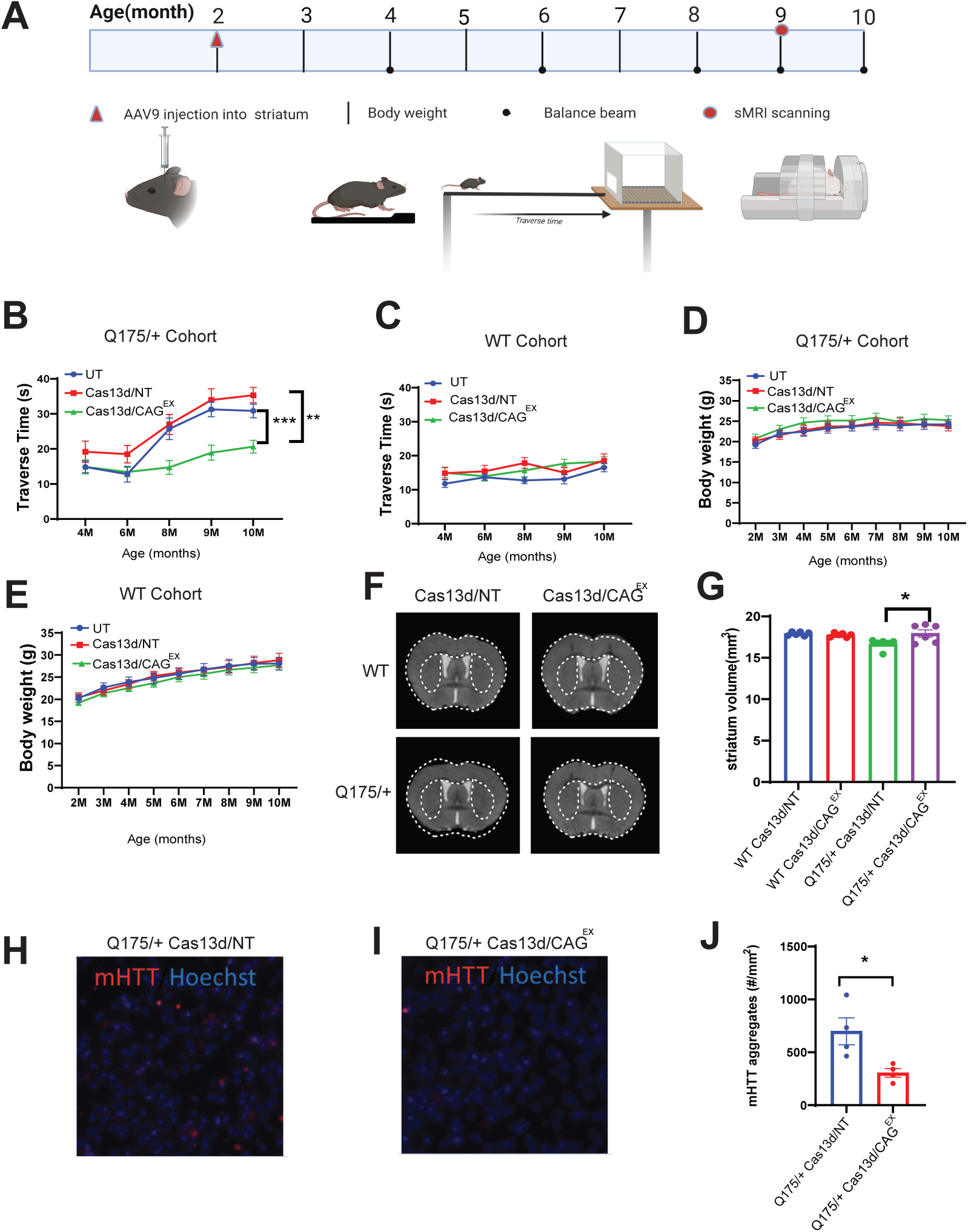
Therapeutic efficacy of Cas13d/CAG^EX^ in a full-length knock-in mouse model of HD. A. Timeline of experimental design and outcome measures. B,C. Mice were tested on a 5 mm balance beam and time crossing the beam (Traverse time) was recorded at indicated ages in zQ175 HD mice or WT mice with indicated treatments. \*\*\**p*<0.001, \*\**p*<0.01, two-way *ANOVA* with Bonferroni post hoc tests. D,E. Body weight in zQ175/+ HD mice or WT mice with indicated treatments. n =10, 5 male and 5 female/group. F. Representative MRI images of the zQ175/+ HD mice and WT mice injected with indicated AAVs. G. Striatal volume was quantified from 3D structural MRI in indicated groups at 9 months of age (7 months after AAV injections). n =6 mice/group, 3 female and 3 male per group. *p<0.05, one-way ANOVA with Bonferroni post hoc analysis. H,I. Representative mHTT aggregates detected by immunostaining with EM48 antibody. Red: EM48 immunostaining; Blue: Hoechst nuclear staining. J. Quantification of mHTT aggregates in the zQ175 mice injected with Cas13d/NT or AAV9-Cas13d/CAG^EX^. n=4. \**p*<0.05 by Student’s t-test.

Next, we examined whether Cas13d/CAG^EX^-mediated mutant *HTT* silencing could improve neuropathology in zQ175/+ mice. Using human-translatable high resolution structural MRI, we delineate regional brain volumes accurately using an automatic segmentation large deformation diffeomorphic metric mapping (LDDMM) including the striatum (Cheng et al 2011, Peng et al 2016, Zhang et al 2010). We performed MRI scans of 9-month-old zQ175/+ mice, when these HD mice display striatal atrophy as we reported previously (Peng et al., 2016). Our behavioral results indicate that Cas13d/NT has no significant effect compared to the untreated control mice, thus we focused on four groups in the MRI study: WT or HD mice injected with Cas13d/NT or Cas13d/CAG^EX^. We demonstrate that zQ175/+ mice injected with Cas13d/NT have significantly reduced striatal volume compared to WT mice with the same injection, while Cas13d/CAG^EX^-injected zQ175/+ mice have preserved striatal volume compared to HD mice injected with Cas13d/NT (Figure 3F-G).

### Allele-selectivity and transcriptome reversal by Cas13d/CAG^EX^ in zQ175/+ mice

As protein aggregates are pathological hallmarks of HD, we determined the effect of AAV9/CAG^EX^ on mutant HTT aggregation in zQ175/+ mouse striatum. Mutant HTT aggregates were immunolabeled by EM48 antibody. Cas13d/CAG^EX^-injected striatum of zQ175/+ mice had significantly reduced EM48^+^ mutant HTT aggregates compared with those injected with Cas13d/NT (Figure 3H-J), indicating that Cas13d/CAG^EX^ can selectively suppress mutant HTT aggregates. Importantly, in addition to phenotypic effects, we observed a significant reduction of mutant HTT protein levels in the striatum of Cas13d/CAG^EX^ -injected zQ175/+ mice. Wildtype HTT protein levels were preserved in both zQ175/+ and WT treated with Cas13d/CAG^EX^, demonstrating allele-specificity of Cas13d/CAG^EX^ (Figure 4A, B, quantified in Figure 4C-G).

**Figure 4.**
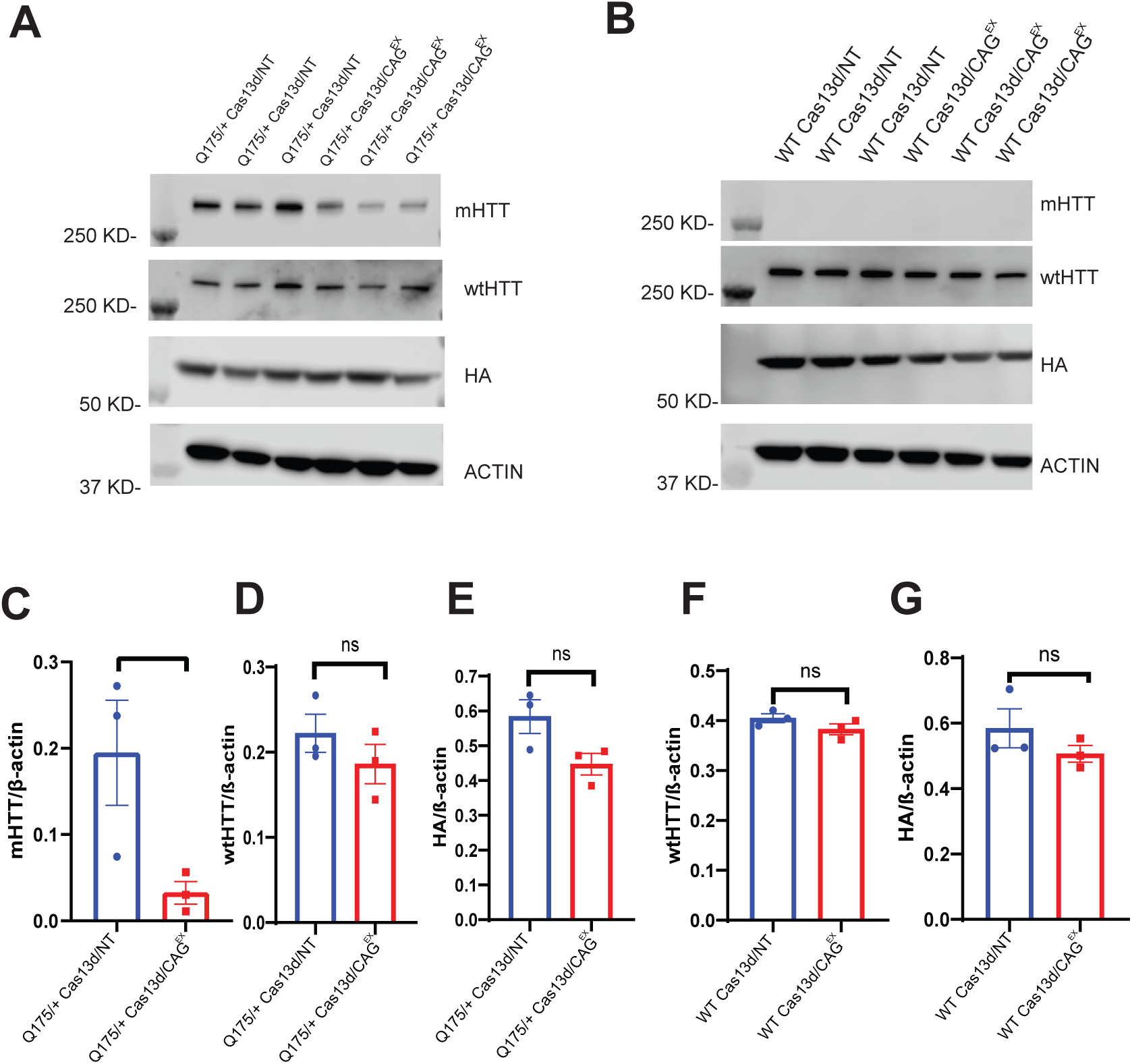
Allele-specific knockdown of mutant HTT protein by Cas13d/CAG^EX^. A, B. Western blot analysis of mutant HTT, wildtype HTT and HA (Cas13d constructs contain HA tag) protein levels in WT and zQ175/+ mice treated with either Cas13d/NT or Cas13d/CAG^EX^. C-G. Quantification of mutant HTT, wildtype HTT and HA protein levels in WT and zQ175/+ mice treated with either Cas13d/NT or Cas13d/CAG^EX^.

At 10 months of age we dissected out the striatum from mice in our cohort and extracted total RNA for RNA-seq analysis. We identified widespread transcriptome dysregulation caused by HD with a total of 1237 significant DEGs in the Cas13d/NT-treated zQ175/+ mice compared to WT littermates (*p*<0.0001, Wilcoxon test, Figure 5A,B, Supplementary Table 2). Gene ontology (GO) analysis showed that genes related to synaptic transmission, dopamine signaling, and behavior were dysregulated in zQ175/+ mice, which is consistent with previous reports (Langfelder et al 2016) (Supplementary Figures 4A, 4B). Hierarchical clustering of the top DEGs showed that the control samples are expectedly distinct from the zQ175/+ samples, but strikingly, zQ175/+ mice treated with Cas13d/CAG^EX^ clustered closer with the WT cohort (Figure 5A). In Cas13d/CAG^EX^-treated zQ175/+ mice, 82.3% of HD-downregulated DEGs and 67.8% of HD-upregulated DEGs had a reduced magnitude fold change relative to WT mice treated with Cas13d/NT, indicating a substantial reversal of HD-associated transcriptome dysregulation (Figure 5C, Supplementary Table 2). Excitingly, as observed with human MSNs, we determined that 64.7% of HD-mediated transcriptional changes of ±2-fold or more were reversed by Cas13d/CAG^EX^ treatment.

**Figure 5.**
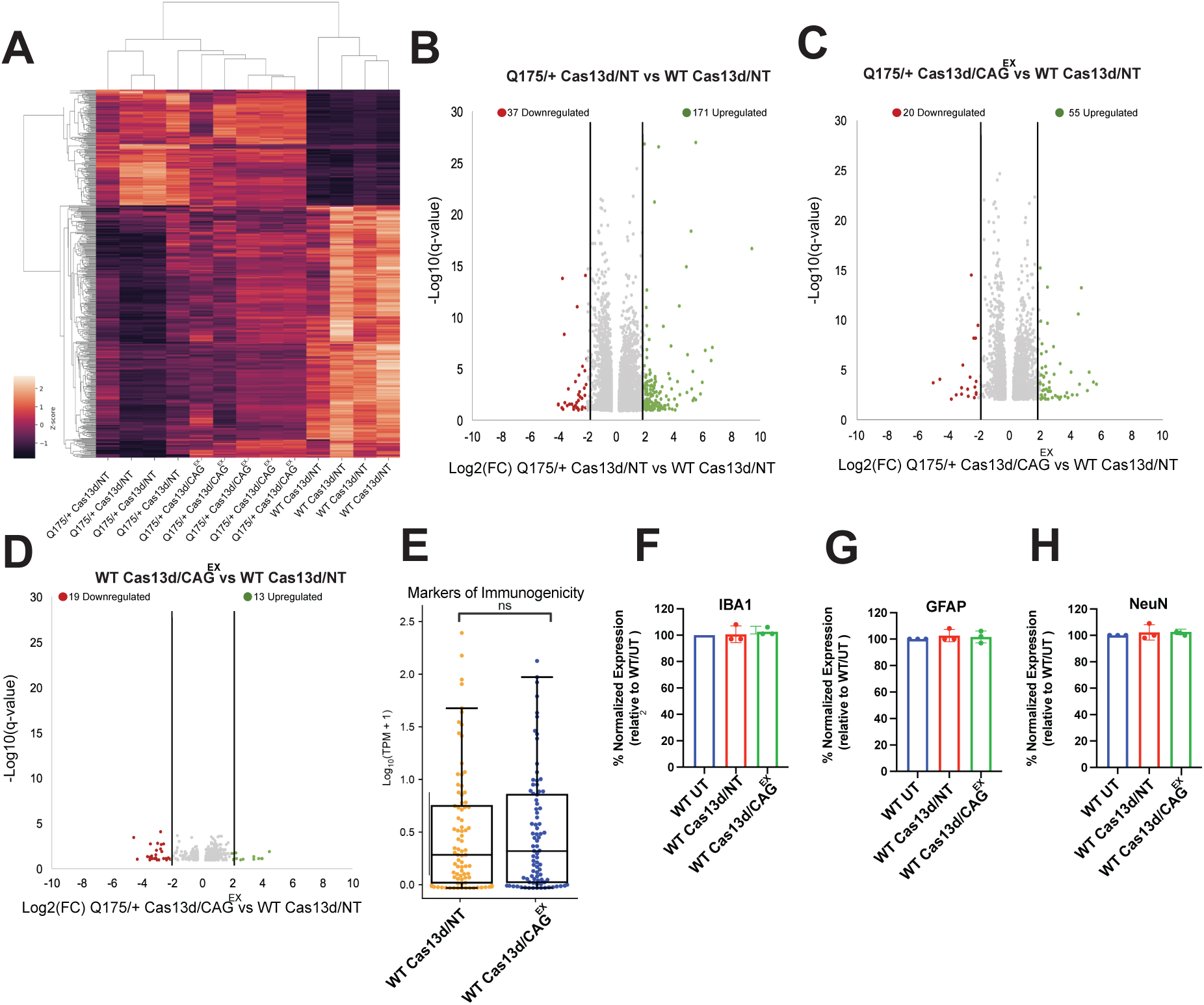
Cas13d/CAG^EX^ partially reverses HD-associated differential gene expression in a full-length knock-in mouse model of HD. Cas13d/CAG^EX^ or Cas13d/NT was injected to the striatum of 2-month-old HD zQ175 or WT mice, and the striatum was harvested at 10 months of age. Total RNA was extracted and quantified by RNA-seq. N = 4 or 5 mice per group. A. Hierarchical clustered heatmap of top 500 differential DEGs in control and HD mouse striatal samples treated with Cas13d/CAG^EX^ or Cad13d/T. B, C Scatter plots of upregulated and downregulated DEGs of within zQ175/+ mice treated with either Cas13d/NT or Cas13d/CAG^EX^ AAV9 showing reversal of HD-mediated transcriptome changes in HD mouse striatum by Cas13d/CAG^EX^ (Wilcoxon Test, p< 0.0001). D. Scatter plots of upregulated and downregulated DEGs of within WT mice treated with either Cas13d/NT or Cas13d/CAG^EX^ AAV9 showing limited off-targets (Wilcoxon Test, p< 0.0001). E. Expression of 100 genes associated with immune system activation (Barta et al 2021). Values reflect average expression of biological replicates for each gene. ns: no significant differences, Wilcoxon test with Benjamini-Hochberg correction. F-H Quantification of CNS-specific markers of immunogenicity via quantitative PCR.

We next evaluated the safety profile of Cas13d/CAG^EX^ based on our RNA-seq data. When comparing WT mice treated with Cas13d/CAG^EX^ or Cas13d/NT, we found only marginal changes in gene expression (Figure 5D, Supplementary Table 2). We next examined the expression levels of 13 genes that contain benign CAG repeats, which could be inadvertently targeted by the CAG^EX^ gRNA. We observed no significant differences in any of these genes (Supplementary Figure 4C). Next, we quantified the expression of 50 immunogenicity-associated genes that were previously reported to be upregulated in preclinical mouse models treated with CRISPR/Cas9-based therapeutics (Batra, 2021). Remarkably, Cas13d/CAG^EX^ treatment did not increase expression of these genes in WT mice, suggesting good tolerance of the treatment by the immune system (Figure 5E). Further, quantitative PCR also confirmed that Cas13d/CAG^EX^ did not disrupt central nervous system-specific markers of immunogenicity including *IBA1*, *GFAP*, and *NeuN* (Figure 5F-H). Taken together, these results support the efficacy and safety of Cas13d/CAG^EX^ in HD preclinical models.

## Discussion

HD is caused by an abnormal expansion of CAG trinucleotide short tandem repeats in exon 1 of the *HTT* gene on chromosome 4 (McColgan et al 2018). Approximately 30,000 people in the United States currently manifest symptoms of HD, but 100,000–200,000 are believed to have inherited this HD causing mutation and would be expected to develop symptoms later in life (Tabrizi et al 2020). Unfortunately, to date, neither curative nor disease-modifying treatment for HD exists. Current therapeutic strategies target different aspects of HD pathology including excitotoxicity, the dopamine pathway, caspases, mHTT protein aggregation, mitochondrial dysfunction, and transcriptional dysregulation (Potkin et al 2018). However, all strategies that target downstream effects of mutant HTT have since failed in human clinical trials. The only therapeutics currently approved by the FDA only alleviate symptoms, and do not modify disease progression.

As the molecular etiology of HD remains complex, it is widely accepted that elimination of the repetitive RNA produced by this locus provides a universal therapeutic principle for patients suffering from HD (Schulte et al 2011). Here, we show extensive preclinical data in two established models of HD for a new gene therapy strategy that effectively eliminates CAG-expanded pre-mRNA with the use of RNA-targeting CRISPR/Cas13d technology. Specifically, we engineered a potent gene therapy vector that encases *Ruminococcus flavefaciens* Cas13d, an RNA-targeting enzyme only 2.4 kb in size, and a CAG-expansion RNA-targeting gRNA in a single viral delivery vehicle. We utilized multiple established preclinical models of HD including the zQ175/+ mouse model which harbors one human allele of HTT with expanded CAG repeats (∼220 repeats) in exon 1, and a series of HD patient iPSC-derived striatal neurons to show both efficacy and safety of our new gene therapy approach both *in vivo* and in a humanized preclinical model.

Numerous strategies exist to attenuate gene expression via targeted destruction of mHTT RNA including antisense oligonucleotides (ASOs), RNA interference compounds, and small molecule splicing modulators. The most promising of these include Roche’s phase III ASO candidate tominersen and Wave Therapeutics’ phase I/II ASO candidates WVE-120101, WVE-120102 (Wild et al 2017). However, each failed to show adequate efficacy and safety in their respective clinical trials. Toxic effects, thought to be caused by either the repeated high dose used or interference with wildtype HTT protein levels, were observed in the case of tominersen which targets both mutant and wildtype HTT pre-mRNA. On the other hand, Wave’s ASO candidates which target patient-specific single nucleotide polymorphisms on mHTT pre-mRNA showed limited target specificity which led to low efficacy. In our study, we show that CAG^EX^ does not disrupt wildtype HTT proteins levels *in vivo* when delivered directly to the striatum of zQ175/+ mice. We also observed its ability to knock down toxic CAG-expanded RNA in a panel of HD patient iPSC-derived striatal neurons, suggesting it could be effective to all patient populations with HD. Both were achieved with limited off-target effects in both the mouse and human striatal neuron transcriptome. This is also an improvement over current RNA-targeting approaches, including small molecules with limited target specificity, as well as RNAi compounds which may deplete endogenous RNAi machinery and cause undesired global alterations in gene expression.

The difficulty associated with delivery to the CNS has thwarted countless therapeutics, but there is a recent resurgence in virally-delivered therapeutics to many tissues including the CNS. Here we show long lasting therapeutic efficacy (8 months post-injection) of CAG^EX^ when delivered by AAV9 *in vivo*. Importantly, AAV9 is an FDA-approved gene therapy delivery method that affords sustained transgene expression and is currently being used in several clinical trials. On the other hand, ASOs and other small molecules must be continually re-administered for the life of the patient, which can pose safety issues in the affected CNS tissues. Taken together, we demonstrate for the first time the proof-of-principle of a CAG^EX^ RNA-targeting CRISPR/Cas13d system as a potential therapeutic approach for HD, a strategy with broad implications for the treatment of other dominantly-inherited neurodegenerative disorders.

## Acknowledgements

We are grateful to the La Jolla Institute’s Immunology Sequencing Core and the IGM Genomics Center, University of California San Diego, for use of the 10X Chromium and Illumina sequencing platforms. This work was supported by National Institutes of Health grants EY029166, NS103172, MH107367, AI132122, HG004659 and HG009889 to G.W.Y.; Bev Hartig Huntington’s Disease Foundation, National Institute of Neurological Disorders and Stroke grants NS099397, NS124084 to W.D. K.H.M was supported by a National Institutes of Neurological Disorders and Stroke NRSA 5 F32 NS112654-03 and a University of California President’s Fellowship. Q.W was a visiting Ph.D student from Beijing University of Chinese Traditional Medicine, PRC (Jan 2019-Dec 2020) and her visit to the Johns Hopkins University School of Medicine was sponsored by a scholarship from the China Scholarship Council and the National Natural Science Foundation of China (No. 82174278, 81973748 to her PhD mentor J.C.).

## Contributions

Conceptualization W.D., K.H.M. and G.W.Y.; Methodology W.D., Q.W., K.H.M. and G.W.Y.; Experimental Work, Q.W., K.H.M., H.L, C.Z, J.C., R.M., K.L., and K.L.J; Formal Analysis, Q.W., K.H.M. and M.L.G; Writing – Original Draft, K.H.M. and Q.W.; Writing – Review & Editing, W.D., K.H.M, and G.W.Y.; Funding Acquisition, W.D and G.W.Y.; Supervision, W.D and G.W.Y.

## Materials and Methods

### Animals

Heterozygous zQ175 (zQ175/+) mice and wild type littermates (both gender- and age-matched in each group) were used in the study. zQ175 line in C57BL/6 background strain was obtained from The Jackson Laboratory (Bar Harbor, ME) and bred and maintained in our lab. Genotyping and CAG repeat size were determined at Laragen Inc. (Culver City, CA, USA) by PCR of tail snips. The CAG repeat length was 223 ± 3 in male zQ175/+ mice and 225 ± 3 in female zQ175 mice/+ used in the study. All mice were housed under specific pathogen-free conditions with a reversed 12-h light/dark cycle maintained at 23C and provided with food and water ad libitum. All behavioral tests and longitudinal MRI measures were done in the mouse dark phase (active). The study was carried out in strict accordance with the recommendations in the Guide for the Care and Use of Laboratory Animals of the National Institutes of Health and approved by Animal Care and Use Committee at Johns Hopkins University. A rigorous study design was implemented. Mice were randomized into groups to avoid bias. Balanced gender ratio was considered in all experiments. Data were collected using animal ID and analyzed by investigators who were blinded to genotype and treatment. Data from different genders were analyzed grouped when there was no gender-dependent difference in the outcome measures.

### Cell Lines

Human iPSC lines derived from individuals with HD and neurotypical controls have been previously characterized elsewhere (Gore, et al; 2011, Smith-Geater et al 2020). iPSC colonies were expanded on Matrigel-coated dishes (BD Biosciences, San Jose, CA, USA) with mTeSR1 medium (StemCell Technologies, Vancouver, Canada). The cells were routinely checked by karyotype and CNV arrays to avoid genomic alterations in the culture. HEK293T cells were purchased from the American Type Culture Collection and were not further authenticated.

### Differentiation of iPSCs into 2D neuronal cultures containing enrichment for medium spiny neurons

Differentiations were performed as follows: iPSC colonies were washed with PBS pH 7.4 (Gibco) and then switched to neural induction medium (Advanced DMEM/F12 (1:1) supplemented with 2 mM GlutamaxTM (Gibco), 2% B27 without vitamin A (Life technologies), 10 μM SB431542, 1 μM LDN 193189 (both Stem Cell Technologies), 1.5 μM IWR1 (Tocris)) with daily medium changes, this was day 0. On day 4 cells were passaged 1:2 with Stempro Accutase (Invitrogen) for 5 minutes at 37C and replated on hESC qualified Matrigel®. At day 8, cells were passaged again 1:2 with Stempro Accutase for 5 minutes at 37C and replated on hESC qualified Matrigel® in a different medium (Advanced DMEM/F12 (1:1) supplemented with 2 mM GlutamaxTM, 2% B27 without vitamin A, 0.2 μM LDN 193189, 1.5 μM IWR1, 20 ng/ml Activin A (Peprotech)). At day 16, cells were plated for neuronal differentiation in SCM1 medium (Advanced DMEM/F12 (1:1) supplemented with 2 mM GlutamaxTM, 2% B27 (Invitrogen), 10 μM DAPT, 10 μM Forskolin, 300 μM GABA, 3 μM CHIR99021, 2 μM PD 0332991 (all Tocris), to 1.8 mM CaCl2, 200 μM ascorbic acid (Sigma-Aldrich), 10 ng/ml BDNF (Peprotech)), medium was 50% changed every 2-3 days. On day 23, there was a full medium change to SCM2 medium (Advanced DMEM/F12 (1:1): Neurobasal A (Gibco) (50:50) supplemented with 2 mM GlutamaxTM), 2% B27, to 1.8 mM CaCl2, 3 μM CHIR99021, 2 μM PD 0332991, 200 μM ascorbic acid, 10 ng/ml BDNF) and medium was 50% changed every 2-3 days until day 32 for harvest (Smith-Geater et al 2020)

### Mycoplasma testing

All iPSC and striatal neuron cultures were routinely tested for mycoplasma by PCR. Media supernatants (with no antibiotics) were collected, centrifuged, and resuspended in saline buffer. Ten microliters of each sample were used for a PCR with the following primers: Forward: GGCGAATGGGTGAGTAAC; Reverse: CGGATAACGCTTGCGACCT. Only negative samples were used in the study.

### Lentivirus Production

Both Cas13/CAG^EX^ and Cas13d/NT constructs were co-transfected with psPAX2 (packaging plasmid) (gift from Didier Trono, Addgene plasmid #12260) and VSV-G (viral envelope) (Stewart et al., 2003) into Lenti-X HEK293T cells (Clontech) at a mass ratio of 5:4:3 using polyethyleneimine (PEI). Viral-containing supernatants were collected and concentrated with Lenti-X Concentrator reagent (Clontech) according to the manufacturer’s protocol (100X concentration) and aliquoted for storage.

### Immunocytochemistry

iPSC-derived striatal neurons were permeabilized with 0.3% Triton X (Sigma-Aldrich) in PBS for 10min and then blocked for 30 minutes in 5% goat serum 2.5% BSA (Gibco) 0.1% Triton X in PBS. Samples were incubated with primary antibodies overnight at 4C (DARPP-32 1:200 Abcam ab40801; FoxP1 1:500 Abcam ab16645; GAD65/67 1:500 Millipore AB1511; MAP-2 1:1000 Millipore MAB3418) and then washed three times with PBS before an overnight 4C incubation in secondary antibody (Alexa-fluor rabbit 555 IgG, mouse 488 IgG (Invitrogen) for 1 hr at 1:500 and subsequently incubated with Hoechst 33342 (Sigma-Aldrich) and mounted with Vectorshield. Samples were imaged with a Zeiss LSM 780 confocal microscope at 10x (Smith-Geater et al 2020).

### Human MSN Bulk RNA-seq library generation

RNA was isolated from ∼100,000 MSNs from 4 independent differentiations per experimental group according to the manufacturer’s protocol and purified using RNeasy columns (Qiagen). A total of 72 mRNA libraries for Illumina sequencing were prepared using KAPA mRNA HyperPrep Kit (KAPA Biosystems, cat. KK8581) from 250 ng of total RNA. Libraries were sequenced on an Illumina Novaseq instrument to a minimum of 3×10^7^ paired-end 100-bp reads.

### Human MSN RNAseq data processing and analysis

RNAseq reads were adapter-trimmed using Cutadapt (v1.14) and mapped to human-specific repetitive elements from RepBase (version 18.05) by STAR (v2.4.0i) (Dobin et al 2013). Repeat-mapping reads were removed, and remaining reads were mapped to the human genome assembly (hg38) with STAR. Read counts for all genes annotated in GENCODE v24 (hg38) were calculated using the read summarization program featureCounts (CDS regions only, Liao et al., 2014). Differential expression analysis between different experimental groups (HD untreated, HD Cas13d/NT, HD Cas13d/CAG^EX^, Control 1-3 untreated, Control 1-3 Cas13d/NT, Control 1-3 Cas13d/CAG^EX^ was performed using DESeq2 (Love et al., 2014). Differentially expressed mRNAs were defined as having a false discovery rate of <0.0001.

### RNA Dot Blot

Human MSN RNA quality and concentrations were estimated using the Nanodrop spectrophotometer. 5 μg of total RNA was resuspended in 1mM EDTA, pH 8.0 to a final volume of 50 μL followed by addition of 30 μL 20X SSC and 20 μL 37% formaldehyde to denature the RNA. The RNA was incubated for 30 min at 60C followed by a 2 minute incubation on ice. The Biorad Bio-Dot apparatus was assembled according to the manufacturer’s protocol with Hybond N+ nylon membrane (GE Healthcare) on the top. The membrane was equilibrated by passing 100 μL of 20X SSC through the slots using vacuum. The denatured RNA was passed through the slots and the membrane was washed with 20X SSC. The membrane was crosslinked at the auto setting in the UV Stratalinker (Stratagene). ULTRAhyb™ Ultrasensitive Hybridization Buffer pre-warmed at 37C was used to pretreat the membrane for 10 min at 37C. A biotinylated probe (CTGCTGCTGCTGCTGCTGCTGCTG-3’biotin (IDT)) was added to the prehybridization solution (5 ul/mL). Hybridization was conducted for 5 hours at 37C and membrane was washed 1X with 1X SSC for 10 minutes at 37C. The membrane was exposed using the Chemiluminescent Nucleic Acid Detection Module Kit (Thermo Fisher).

### AAV packaging

Cas13d/CAG^EX^ (#1457 Cas13d/CAG^EX^: 3.9×10^12^ vg/ml) and Cas13d/NT (#1457 Cas13d/NT: 4.4 × 10^12^ vg/ml) were cloned into an AAV vector provided by UCSD Viral Vector Core. Each vector was packaged into single-stranded AAV9 by both the UCSD Viral Vector Core (UCSD, La Jolla, CA) and the Emory Viral Vector Core (Emory University, Atlanta, GA) using standard procedures and quality controls established at each independent core.

### Stereotaxic injection

The mice were anesthetized with 1.5% isoflurane inhalation and stabilized in a stereotaxic instrument (David Kopf Instruments). Mice were injected into the striatum using the stereotaxic coordinates: 0.62mm rostral to bregma, ±1.75 mm lateral to midline and 3.5 mm ventral to the skull surface. 1 μl of a Cas13d/CAG^EX^ – containing AAV9 titer (3.9 ×10^12^ vg/ml) or Cas13d/NT – containing AAV9 titer (4.4 ×10 ^12^ vg/ml, diluted to 3.9 ×10^12^ vg/ml) were injected into the striatum using a Hamilton syringe infusion pump (World Precision Instruments, Sarasota, Florida, USA) at a perfusion speed of 200 nl/min.

### Immunohistochemistry and Quantification

Mice were anesthetized and perfused transcardially with phosphate-buffered saline (PBS) followed by 4% paraformaldehyde. Brains were post-fixed overnight followed by immersion in 30% sucrose for 24 h. Frozen sections (thickness, 10-15 μm) of coronal brain were immunostained with following antibodies, EM48 (MAB5347, 1:200, Merck Millipore, USA). Briefly, the sections were washed three times with PBS, then permeabilized by incubating with 0.3% Triton X-100 for 5 min, followed by incubation with blocking solution containing 5% donkey serum or 3% goat serum and 0.3% Triton X-100 for 1 h. The sections were then incubated with primary antibody at 4C overnight. After three washings with PBS, the sections were incubated with fluorescence-labeled secondary antibody for 2 h at room temperature. Sections were mounted onto Superfrost slides (Fisher Scientific, Pittsburgh, PA, USA) dried and then covered with anti-fade mounting solution. Fluorescence images were acquired with Keyence BZ-X700 All-in-One florescence microscope.

For image analysis, the samples were coded with ID and images were analyzed by investigators who are blinded to the genotypes and treatment. Analysis results were then calculated statistically and decoded by different investigators at the end. The results from three microscopy fields per slide and three sections per mouse were calculated for each mouse brain. EM48 positive puncta quantities were determined using the Fuji-ImageJ’s analyze particles plugin function. The numbers of EM48 positive puncta per mm^2^ were determined at a constant threshold for each stain using × 20 images for quantifications

### Western Blotting

Striatal samples were homogenized in a buffer containing 50 mM Tris-HCl (pH 8.0), 150 mM NaCl, 0.1% (w/v) SDS, 1.0% NP-40, 0.5% sodium deoxycholate, and 1% (v/v) protease inhibitors. 30 μg of proteins were separated in a 4-20% gradient gel and transferred to a nitrocellulose membrane. The membrane was blotted with the following primary antibodies: MAB 2166 (anti-HTT, 1:1000, Millipore Sigma, USA), MW1 (anti-poly-Q, 1:1000, Millipore Sigma, USA), and β-actin (Sigma, mouse monoclonal antibody, 1:5000). After incubation with HRP-conjugated secondary antibodies, bound antibodies were visualized by chemiluminescence.

### *In Vivo* structural MRI acquisition

*In vivo* MRI was performed on a vertical 9.4 Tesla MR scanner (Bruker Biospin, Billerica, MA, USA) with a triple-axis gradient and a physiological monitoring system (EKG, respiration, and body temperature). Mice were anesthetized with isoflurane (1%) mixed with oxygen and air at 1:3 ratios via a vaporizer and a facial mask and scanned longitudinally (the same mice were imaged repeatedly over a 12-month period). We used a 20-mm diameter volume coil as the radiofrequency transmitter and receiver. Temperature was maintained by a heating block built into the gradient system. Respiration was monitored throughout the entire scan.

High-resolution anatomical images were acquired by using a three-dimensional (3D) T2-weighted fast spin echo sequence with the following parameters: echo time (TE)/repetition time (TR) = 40/700 ms, resolution = 0.1 mm × 0.1 mm × 0.25 mm, echo train length = 4, number of average = 2, and flip angle = 40°. Multi-slice T2-weighted images of the mouse brain were acquired by the RARE (Rapid Acquisition with Refocused Echoes) sequence with the following parameter (echo time (TE) / repetition time (TR) = 40 ms/1500 ms, RARE factor = 8, in-plane resolution = 0.125 mm x 0.125 mm, slice thickness = 1 mm, total imaging time less than 2 min) and used for high resolution anatomical imaging. Total imaging time was about 50 min per mouse. Mice recovered quickly once the anesthesia was turned off, and all mice survived the imaging sessions.

### Structural MRI image analysis

Images were first rigidly aligned to a template image by using automated image registration software (http://bishopw.loni.ucla.edu/AIR5/, AIR). The template image was selected from one of the images acquired from age-matched littermate control mice (mouse had the medium brain volume among the control group), which had been manually adjusted to the orientation defined by the Paxinos atlas with an isotropic resolution of 0.1 mm x 0.1 mm x 0.1 mm per pixel. After rigid alignment, images had the same position and orientation as the template image, and image resolution was also adjusted to an isotropic resolution of 0.1 mm × 0.1 mm × 0.1 mm per pixel.

Signals from non-brain tissue were removed manually (skull-stripping). Skull-stripped, rigidly aligned images were analyzed by using Landmarker software (www.mristudio.org). Intensity values of the gray matter, white matter, and cerebral spinal fluid were normalized to the values in the template images by using a piece-wise linear function. This procedure ensured that subject image and template image have similar intensity histograms. The intensity-normalized images were submitted by Landmarker software to a linux cluster, which runs Large Deformation Diffeomorphic Metric Mapping (LDDMM). The transformations were then used for quantitative measurement of changes in local tissue volume among different mouse brains, by computing the Jacobian values of the transformations generated by LDDMM. There are 29 different brain regions segmented automatically.

### Behavioral tests

5mm balance beam testing was conducted on an 80-cm long and 5-mm wide square shaped balance beam that was mounted on supports of 50-cm in height. A bright light illuminated the start platform, and a darkened enclosed 1728 cm^3^ escape box (12 × 12 × 12 cm^3^) was situated at the end of the beam. Mice were trained to walk across the beam twice at least 1 h prior to testing. The time for each mouse to traverse the balance beam was recorded with a 125-sec maximum cut-off, and falls were scored as 125 sec.

### Mouse RNA isolation and RNA-seq library preparation

RNA was isolated from snap frozen striatal mouse tissue from 2 independent cohorts of mice according to the manufacturer’s protocol and purified using RNeasy columns (Qiagen). A total of 13 mRNA libraries for Illumina sequencing were prepared using KAPA mRNA HyperPrep Kit (KAPA Biosystems, cat. KK8581) from 250 ng of total RNA. Libraries were sequenced on an Illumina Novaseq instrument to a minimum of 2×10^7^ paired-end 100-bp reads.

### Mouse RNA-seq data processing and analysis

To correct for batch effects between library preparations, ComBat-seq was applied to the raw read counts (Zhang et al., 2020). RNA-seq reads were adapter-trimmed using Cutadapt (v1.14) and mapped to mouse-specific repetitive elements from RepBase (version 18.05) by STAR (v2.4.0i) (Dobin et al., 2013). Repeat-mapping reads were removed, and remaining reads were mapped to the mouse genome assembly (mm10) with STAR. Read counts for all protein-coding genes annotated in GENCODE (vM20) were calculated using the read summarization program featureCounts (Liao et al., 2014). Differential expression analysis was performed using DESeq2 (Love et al., 2014). Differentially expressed mRNAs were defined as having a false discovery rate of <0.01.

### Human and mouse gene ontology analysis

PANTHER Version 16 was used for GO analysis (Mi et al 2013). All human or mouse protein-coding genes with at least 5 reads in any sample were used as the background set. Significant GO terms were determined by Fisher’s exact test after FDR correction at P < 0.05 and sorted by fold enrichment.

### Quantitative PCR

For qPCR, RNA was converted to cDNA using the High-Capacity cDNA Reverse Transcription Kit (ThermoFisher). Quantification of mRNA in both human cells and mouse tissue samples were completed with PrimeTime Gene Expression Master Mix (IDT) and IDT Prime-time qPCR assays: (Grik2 Hs.PT.58.39107351 & Mm.PT.58.16642818; Darpp2 Hs.PT.58.40063116 & Mm.PT.58.9253526); Pde10a (Hs.PT.58.430241 & Mm.PT.58.16919824). Human and Mouse Actn2 IDT Prime-time qPCR assays (Hs.PT.56a.39451065 & Mm.PT.58.28953379) were used a an internal, housekeeping control.

### Statistical Analysis

Data are expressed as the mean ± SEM unless otherwise noted. Student’s *t*-test was used to measure the significant levels between WT and zQ175/+ HD groups at each given age. Two-way repeated *ANOVA* (genotype and treatment) was performed to examine the statistical difference between different groups along with age and treatment. Statistical analysis was performed with GraphPad software. The *p*-values less than 0.05 were considered as statistically significant. N is reported in the figure legends.

### Data Availability

All the data supporting the findings of this study are available upon reasonable request.

---

**Supplementary Figure 1.**
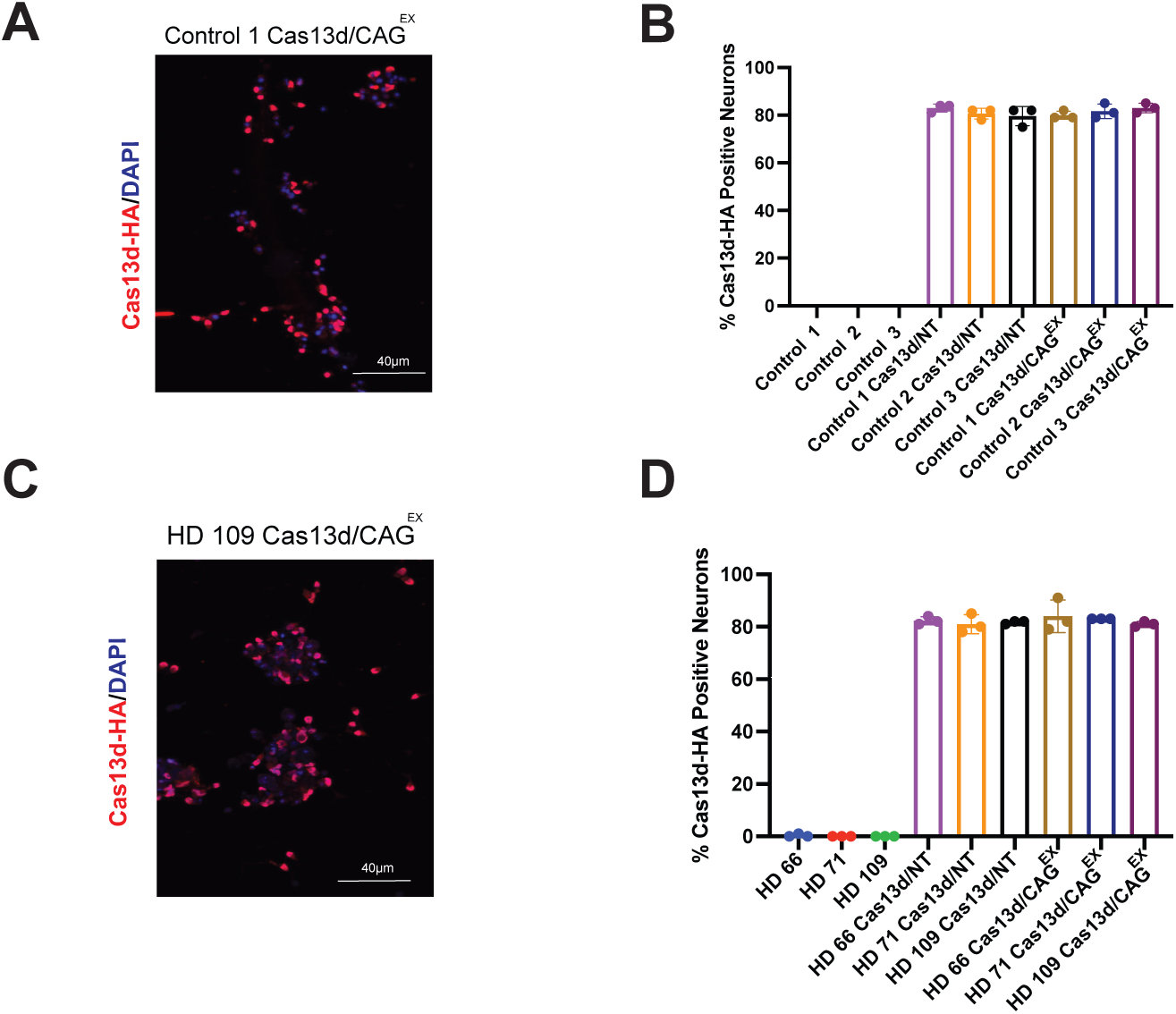
Cas13d Distribution in iPSC-derived MSNs. A-D Image and Quantification of Cas13d-HA distribution in Control and HD Day 32 MSN cultures.

**Supplementary Figure 2.**
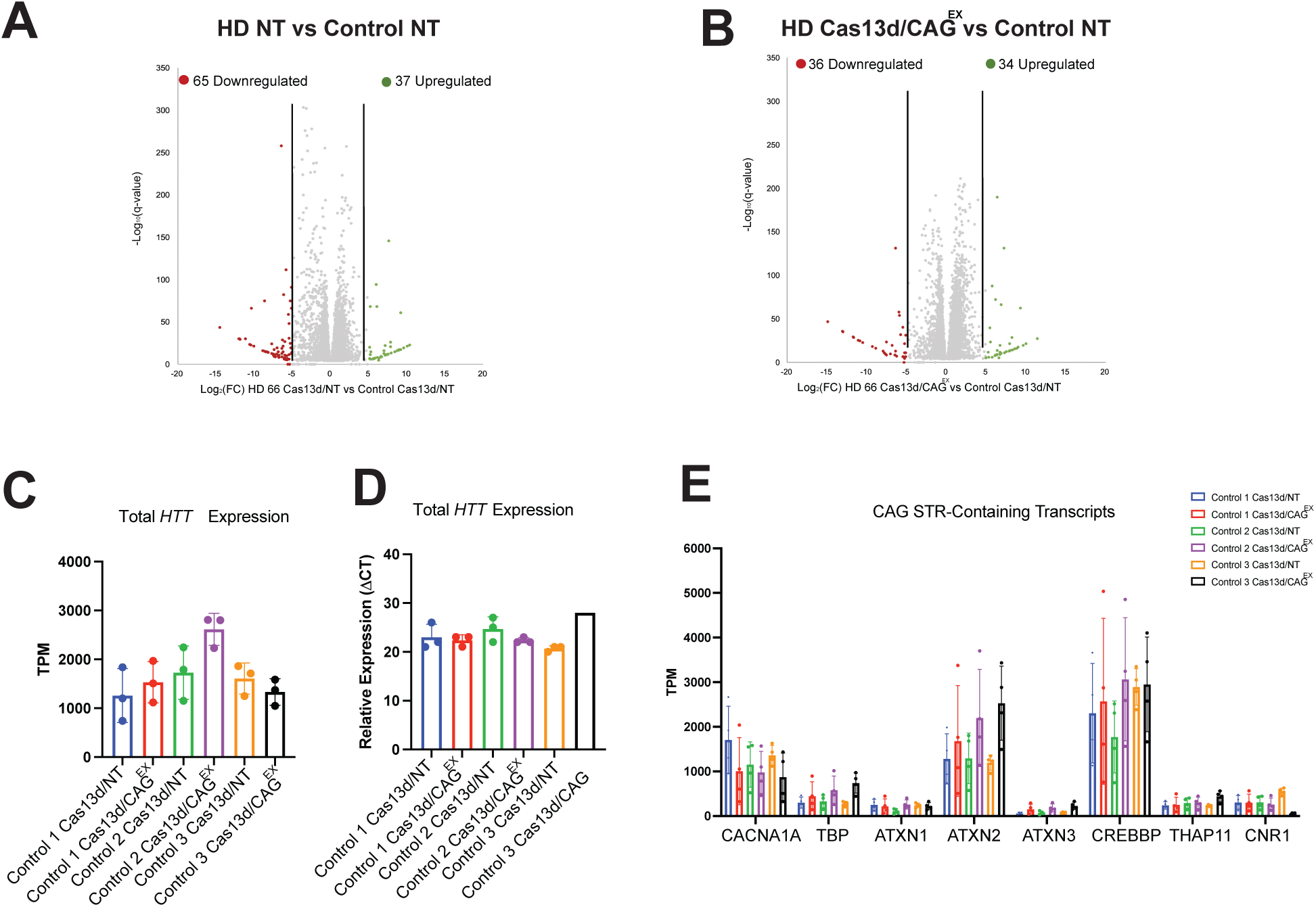
Allele-specificity and safety of Cas13d/CAG^EX^ in a Full-length mHTT Knock-in Mouse Model. A,B. Scatter plots of upregulated and downregulated DEGs of within the striatum of control mice or zQ175 HD mice treated with either Cas13d/NT or Cas13d/CAG^EX^ AAV9 showing reversal of HD-mediated changes in the transcriptome by Cas13d/CAG^EX^ (Wilcoxon Test, p< 0.0001). C,D. Quantification of total *HTT* transcript by TPM and quantitative PCR within Control mouse striatum treated with Cas13d/NT or Cas13d/CAG^EX^. E. Quantification of 8 transcripts with known CAG STRs by TPM in Control mice treated with Cas13d/NT or Cas13d/CAG^EX^.

**Supplementary Figure 3.**
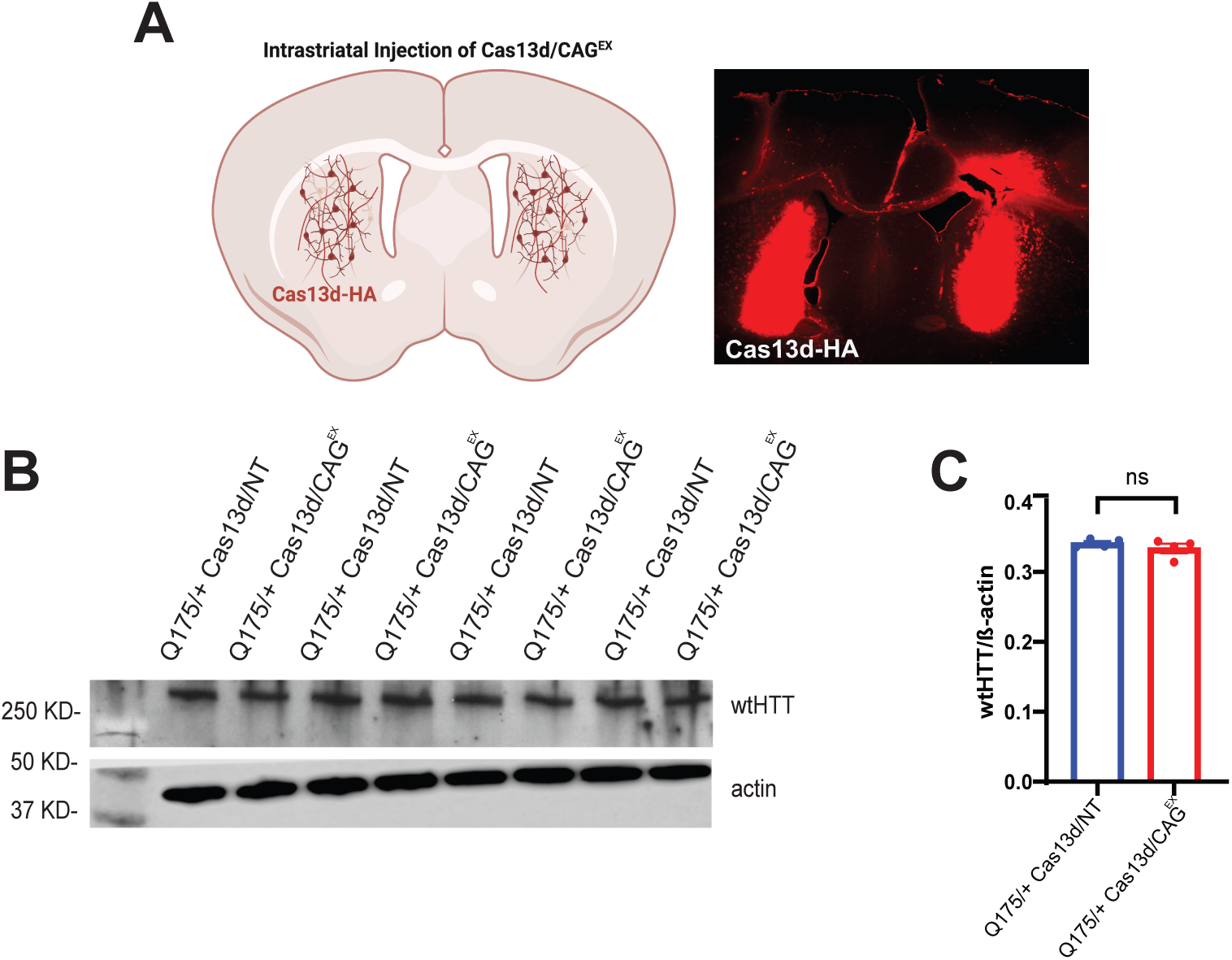
Allele-specificity and safety of Cas13d/CAG^EX^ in a Full-length mHTT Knock-in Mouse Model. A Detection of by Cas13d-HA via HA immunostaining (red) 3 weeks post-intrastriatal injection of AAV9-Cas13d/CAG^EX^. B, C. Western blot analysis of wild type HTT (wtHTT, antibody MAB 2166 antibody). There are no significant differences between two groups by Student’s *t*-test.

**Supplementary Figure 4.**
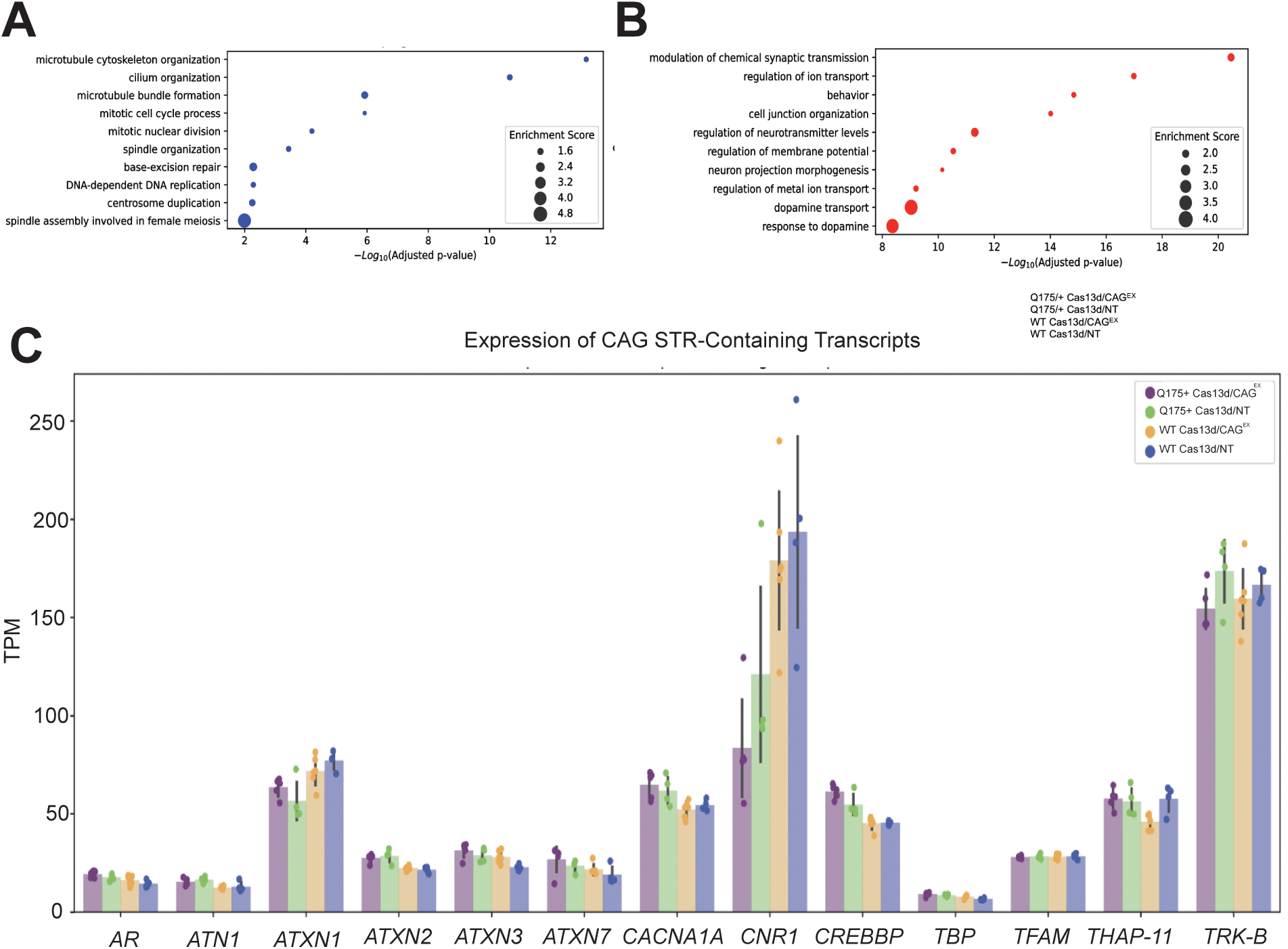
A, B GO analysis of top 500 upregulated and downregulated DEGs in zQ175/+ C. Expression of 13 genes containing CAG repeats. There were no significant differences between treatment groups for any gene (2-sided Manning-Whitney-U test).

