## Supplementary material for "RNA-Targeting CRISPR/Cas13d System Eliminates Disease-Related Phenotypes in Pre-clinical Models of Huntington’s Disease": Methods & Supplement Materials

**Data Availability**

All the data supporting the findings of this study are available upon reasonable request.

**Supplementary Figure 1.**

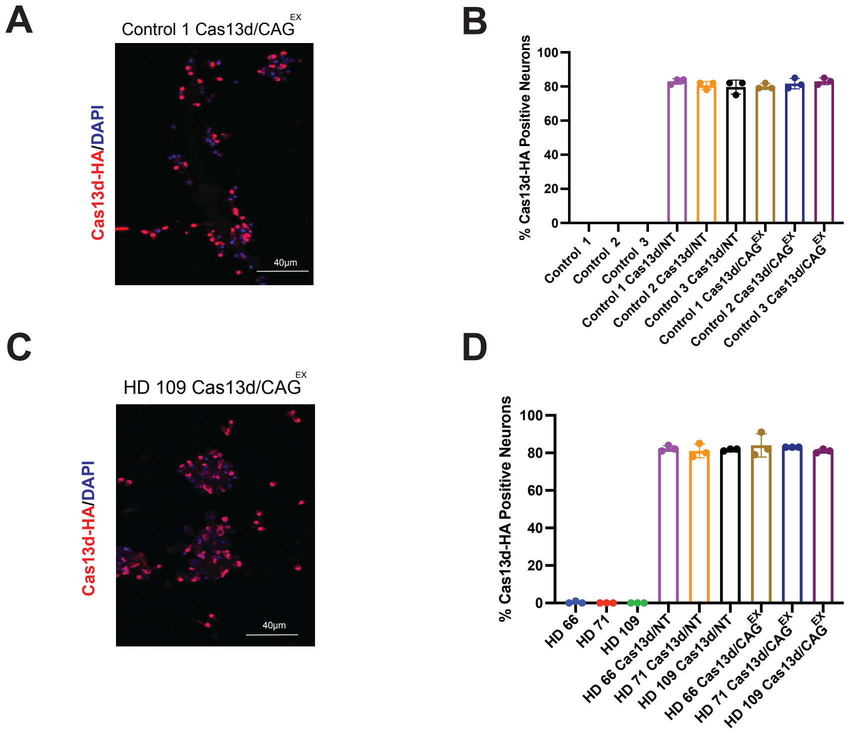


.

**Supplementary Figure 1. Cas13d Distribution in iPSC-derived MSNs**

A-D Image and Quantification of Cas13d-HA distribution in Control and HD Day 32 MSN cultures.

**Supplementary Figure 2.**

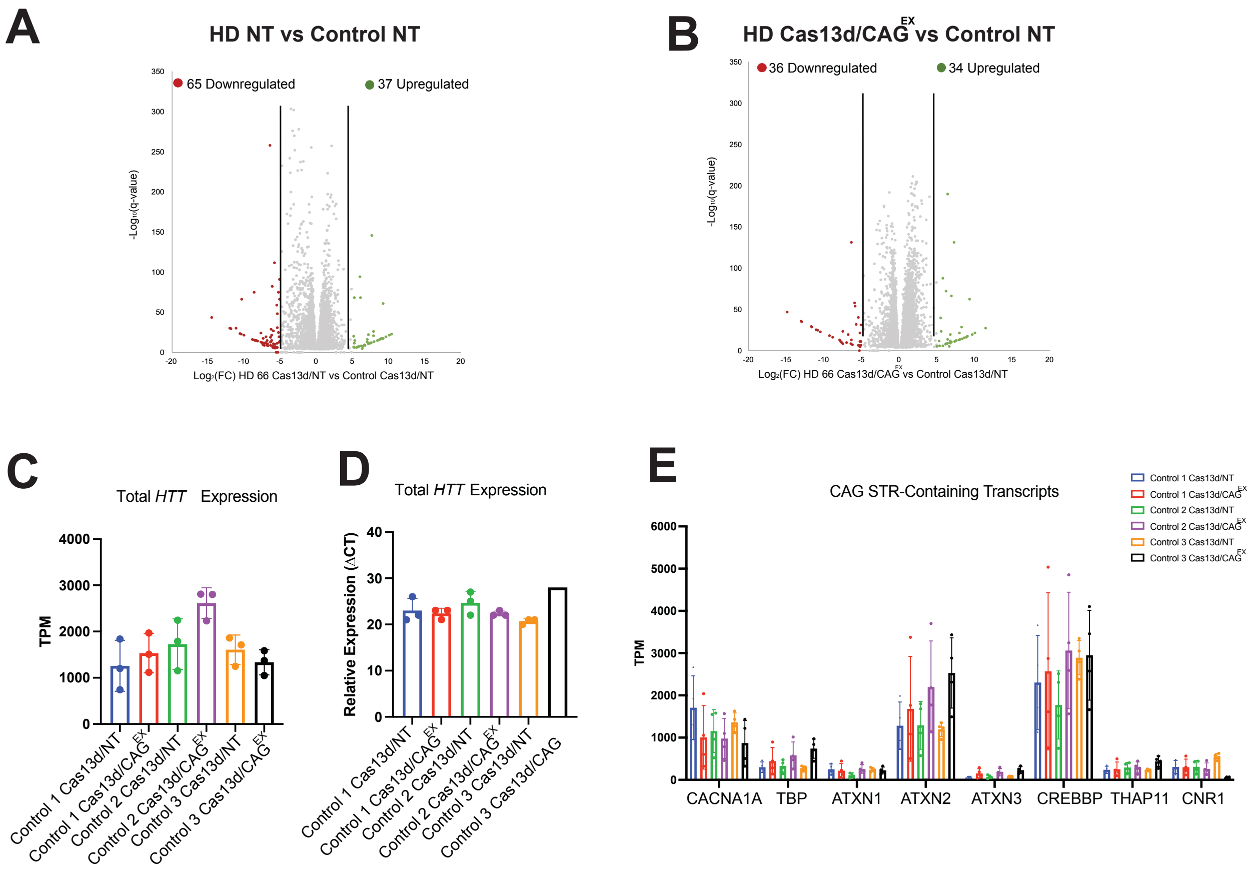

**Supplementary Figure 2. Allele-specificity and safety of Cas13d/CAG$^{EX}$ in a Full-length mHTT Knock-in Mouse Model.**

A,B. Scatter plots of upregulated and downregulated DEGs of within the striatum of control mice or zQ175 HD mice treated with either Cas13d/NT or Cas13d/CAG$^{EX}$ AAV9 showing reversal of HD-mediated changes in the transcriptome by Cas13d/CAG$^{EX}$ (Wilcoxon Test, $p< 0.0001$).

C,D. Quantification of total *HTT* transcript by TPM and quantitative PCR within Control mouse striatum treated with Cas13d/NT or Cas13d/CAG$^{EX}$.

E. Quantification of 8 transcripts with known CAG STRs by TPM in Control mice treated with Cas13d/NT or Cas13d/CAG$^{EX}$.

### Supplementary Figure 3.

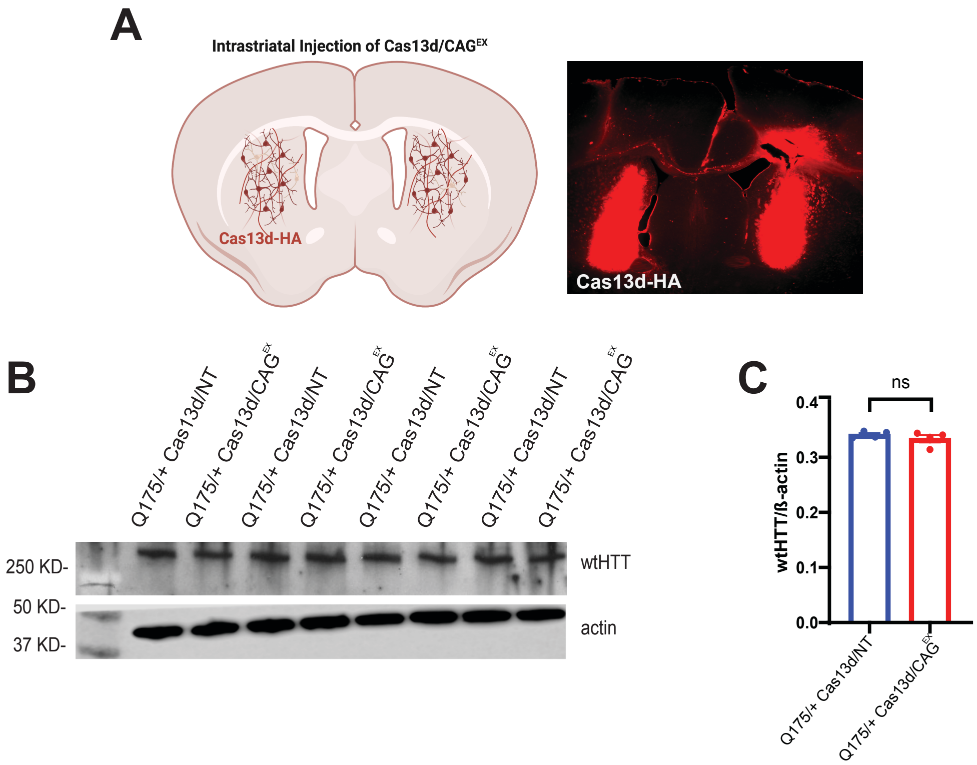

**Supplementary Figure 3. Allele-specificity and safety of Cas13d/$CAG^{EX}$ in a Full-length mHTT Knock-in Mouse Model.**

A Detection of by Cas13d-HA via HA immunostaining (red) 3 weeks post-intrastriatal injection of AAV9-Cas13d/$CAG^{EX}$.

B, C. Western blot analysis of wild type HTT (wtHTT, antibody MAB 2166 antibody). There are no significant differences between two groups by Student's *t*-test.

### Supplementary Figure 4.

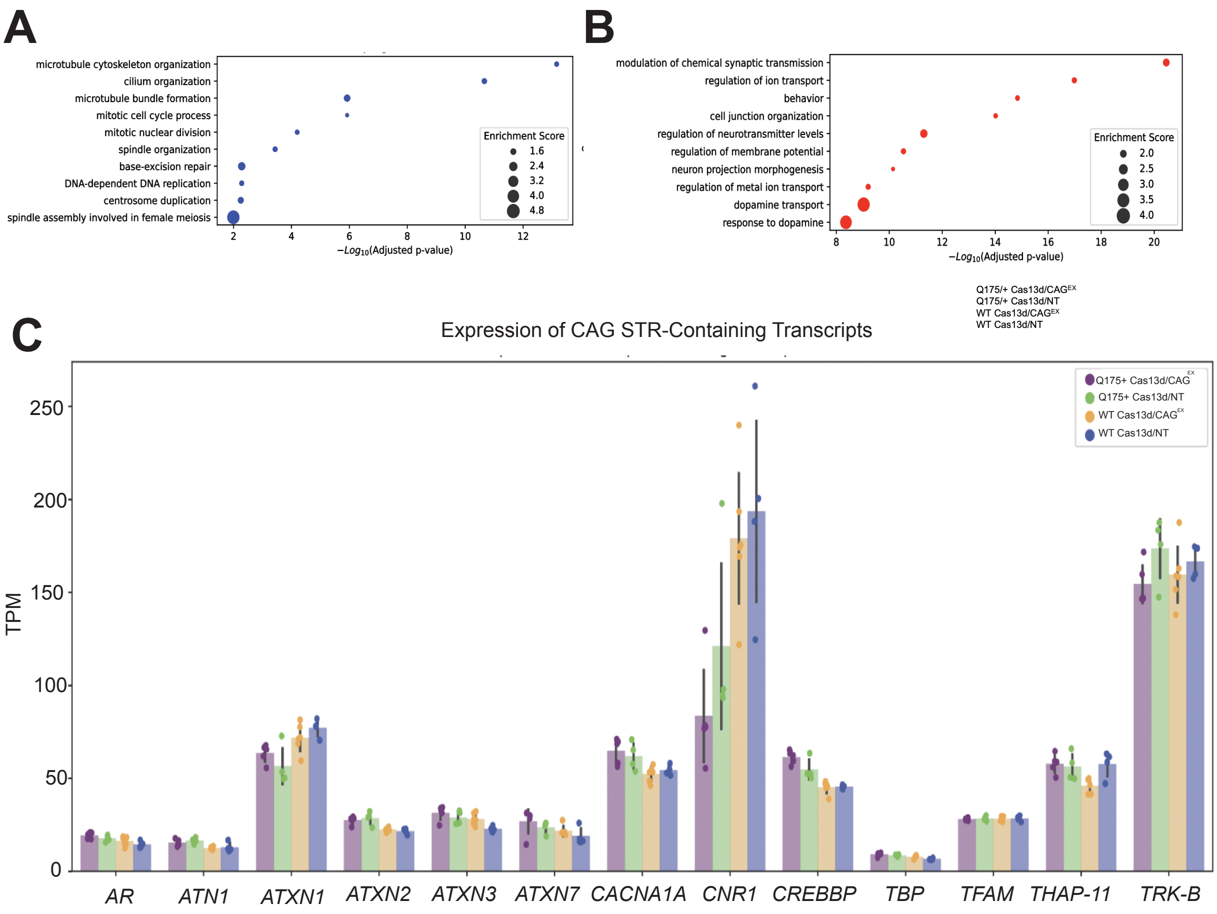

**Supplementary Figure 4.**

A, B GO analysis of top 500 upregulated and downregulated DEGs in zQ175/+

C. Expression of 13 genes containing CAG repeats. There were no significant differences between treatment groups for any gene (2-sided Manning-Whitney-U test).
